# The Hsc70 Disaggregation Machinery Removes Monomer Units Directly from α-Synuclein Fibril Ends

**DOI:** 10.1101/2020.11.02.365825

**Authors:** Matthias M. Schneider, Saurabh Gautam, Therese W. Herling, Ewa Andrzejewska, Georg Krainer, Alyssa M. Miller, Quentin A. E. Peter, Francesco Simone Ruggeri, Michele Vendruscolo, Andreas Bracher, Christopher M. Dobson, F. Ulrich Hartl, Tuomas P. J. Knowles

## Abstract

Molecular chaperones contribute to the maintenance of cellular protein homeostasis through a wide range of mechanisms, including the assistance of *de novo* protein folding, the rescue of misfolded proteins, and the prevention of amyloid formation. Chaperones of the Hsp70 family have a striking capability of disaggregating otherwise irreversible aggregate structures such as amyloid fibrils that accumulate during the development of neurodegenerative diseases. However, the mechanisms of this key emerging functionality remain largely unknown. Here, we bring together microfluidic measurements with kinetic analysis and show that that the Hsp70 protein heat chock complement Hsc70 together with its two co-chaperones DnaJB1 and the nucleotide exchange factor Apg2 is able to completely reverse the aggregation process of alpha-synuclein, associated with Parkinson’s disease, back to its soluble monomeric state. Moreover, we show that this reaction proceeds with first order kinetics in a process where monomer units are taken off directly from the fibril ends. Our results demonstrate that all components of the chaperone triad are essential for fibril disaggregation. Lastly, we quantify the interactions between the three chaperones as well as between the chaperones and the fibrils in solution, yielding both binding stoichiometries and dissociation constants. Crucially, we find that the stoichiometry of Hsc70 binding to fibrils suggests Hsc70 clustering at the fibril ends. Taken together, our results show that the mechanism of action of the Hsc70–DnaJB1–Apg2 chaperone system in disaggregating α-synuclein fibrils involves the removal of monomer units without any intermediate fragmentation steps. These findings are fundamental to our understanding of the suppression of amyloid proliferation early in life and the natural clearance mechanisms of fibrillar deposits in Parkinson’s disease, and inform on the possibilities and limitations of this strategy in the development of therapeutics against synucleinopathies and related neurodegenerative diseases.

## Introduction

Misfolding and aggregation of proteins and peptides into amyloidogenic fibrils are hallmarks of a wide range of neurodegenerative disorders^1–3^, including α-synuclein (αS) in Parkinson’s disease, the Aβ-peptide in Alzheimer’s disease, and Huntingtin (HTT) in Huntington’s disease^4^. The accumulation of such fibrillar deposits in the central nervous system occurs in an age-dependent manner; earlier in life this process is counteracted by efficient cellular protein quality control machineries that inhibit amyloid formation and thus disease onset earlier in life^5–7^.

Molecular chaperones are critical components of this quality control system^7^. Initially identified as part of the heat stress response^8–10^, chaperones have been shown to assist protein folding and rescue misfolded states^6,8^. Particularly variants of the 70 kDa heat shock protein family (Hsp70s) populate some of the most critical nodes in the proteostasis network and are involved in assisting protein folding, in exerting holdase activity, in preventing protein aggregation, and in degrading misfolded proteins, as well as in mediating the assembly and disassembly of oligomeric protein species^7,11,12^. Hsp70s along with various other chaperones, have been shown to modulate essentially all microscopic steps in amyloid formation, including elongation^13,14^, primary nucleation^15^, as well as secondary nucleation^16^. In recent years, mounting evidence has indicated that Hsp70 chaperones are also involved in the disassembly of aggregates and capable of disaggregating even persistent amyloidogenic aggregate structures such as αS, HTT and Tau fibrils^17–26^. In particular, the constitutively expressed chaperone heat shock complement Hsc70 (HSPA8) together with the Hsp40 class B J-protein DnaJB1 and the Hsp110 family nucleotide exchange factors (NEFs) Apg2 or Hsp105α have been demonstrated to constitute a powerful ATP-driven disaggregase system that disassembles amyloids within minutes via combined fibril fragmentation into shorter fibrils, promoting their depolymerisation into monomers or smaller oligomeric structures^17–19^.

Several studies have provided substantial insights in the basic working principles of the Hsc70–DnaJB1–Hsp110 triad chaperone system (Fig. 1a)^7,18,27^. Structurally, Hsc70, as other Hsp70 chaperones, contains an N-terminal nucleotide-binding domain of 40 kDa, which is linked via a flexible, hydrophobic linker to a 15-kDa substrate-binding domain (SBD) and a 10-kDa α-helical lid^7,11,12^. The SBD recognises hydrophobic peptide segments that are exposed in non-native substrate proteins^28–30^, which are delivered to Hsp70 by J-protein chaperones such as DnaJB1. Upon ATP hydrolysis to ADP induced by the J-protein, the lid closes to form a stable substrate–Hsp70 complex and the J-protein dissociates from Hsp70^7,11,12^. NEFs, in particular members of the Hsp110 family (e.g., Apg2 or Hsp105α)^31–35^, replace the bound ADP with ATP, facilitating lid-opening and substrate release. Since assembly of the chaperone machinery on a protein aggregate leads to a significant entropy loss due to excluded volume effects, the chaperone is thought to act on its substrate by entropic pulling, that is, by exerting a force of up to 15–20 pN to the region it is bound to, leading to the fibril disaggregation^36–38^.

**Figure 1:**
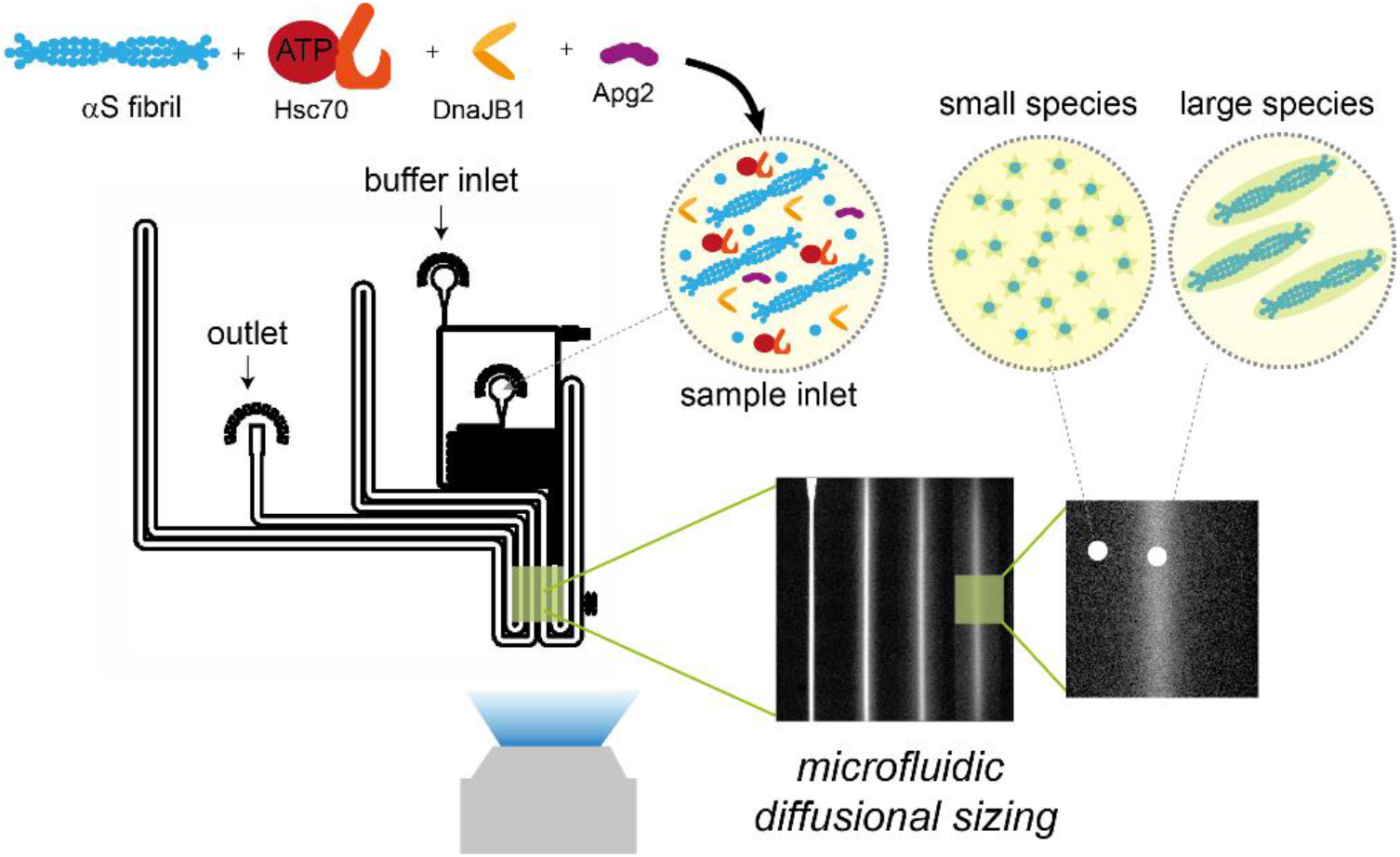
Hsc70-mediated αS fibril disaggregation monitored by microfluidic diffusional sizing. αS fibrils are incubated with the Hsc70–DnaJB1–Apg2 chaperone triad. At different time points, the reaction mixture is analysed using microfluidic diffusional sizing. From the recorded diffusion profiles, the size decay with time is monitored, allowing kinetic and mechanistic analysis of the disaggregation reaction.

While basic aspects of the functional cooperation of the Hsc70–DnaJB1–Apg2 chaperone system in protein disaggregation have been established,^7,18^ key mechanistic questions remain unanswered. Specifically, it is unclear if the disaggregase machinery acts on the fibril ends or along the fibril surface, and if monomer is removed from the fibril substrate, or whether smaller fragments are produced. Moreover, fundamental functional and biophysical parameters such as binding stoichiometry and binding affinities describing the interaction between the participating components are yet to be resolved. Insights into these aspects are of fundamental importance, not least because of the therapeutic potential of Hsp70-mediated disaggregation in neurodegeneration.

Here, we dissect the molecular mechanisms by which the Hsc70–DnaJB1–Apg2 chaperone system disaggregates αS fibrils. We employ microfluidic diffusional sizing in conjunction with chemical kinetics analysis to quantify and characterise the molecular species formed during disaggregation (Fig. 1). A key finding of this study is that the chaperones disassemble αS fibrils into monomers. We show that this process follows pseudo-first order kinetics, which suggests that monomer units are removed directly from the fibril ends. This finding is supported by results obtained from single-round disaggregation experiments, which clearly show occurrence of monomer after a single disaggregation round. All components of the disaggregation system, including Hsc70, Apg2, DnaJB1 and ATP, are essential for the disaggregation reaction, as disassembly does not proceed in absence of either of them. Lastly, we assess the binding properties between the different chaperones and co-chaperones as well as between the chaperones and the fibrils. Based on these results, we establish, for the first time, a full kinetic and thermodynamic profile of the Hsc70-mediated disaggregation reaction and propose a model of αS disaggregation suggesting that Hsc70 chaperones form a cluster in order to exhibit disaggregase functionality on αS fibrils.

## Results

### Kinetics of αS Fibril Disaggregation by the Hsc70 Chaperone Machinery

We applied a microfluidic diffusional sizing approach^39–41^ to investigate the time evolution of αS fibril disaggregation by the Hsc70 chaperone machinery. Such heterogeneous multi-component systems can be challenging to study using conventional surface-based analysis approaches, but the absence of convective mixing on microfluidic scales allows these interactions to be investigated directly in solution without the requirement for any of the binding partners to be immobilised onto a surface. The principle of the technique consists in measuring the molecular diffusivity of protein components and monitoring changes as they undergo binding events. In practice, we capture the diffusion process in both space and time by acquiring the longitudinal diffusion profiles of protein molecules, here αS species, flowing in a microfluidic channel (Fig. 2f). The diffusion profiles acquired in these experiments are then analysed by considering advection–diffusion processes to extract the distribution of diffusion coefficients (not shape-dependent) and the corresponding hydrodynamic radii (*R*_h_) of the individual species present in solution^41^. Smaller species result in a broader profile, whereas larger species remain at the centre of the channel. Crucially, this method also allows for the detection of protein–protein interactions by monitoring the increase in size associated with binding^16,42^.

**Figure 2:**
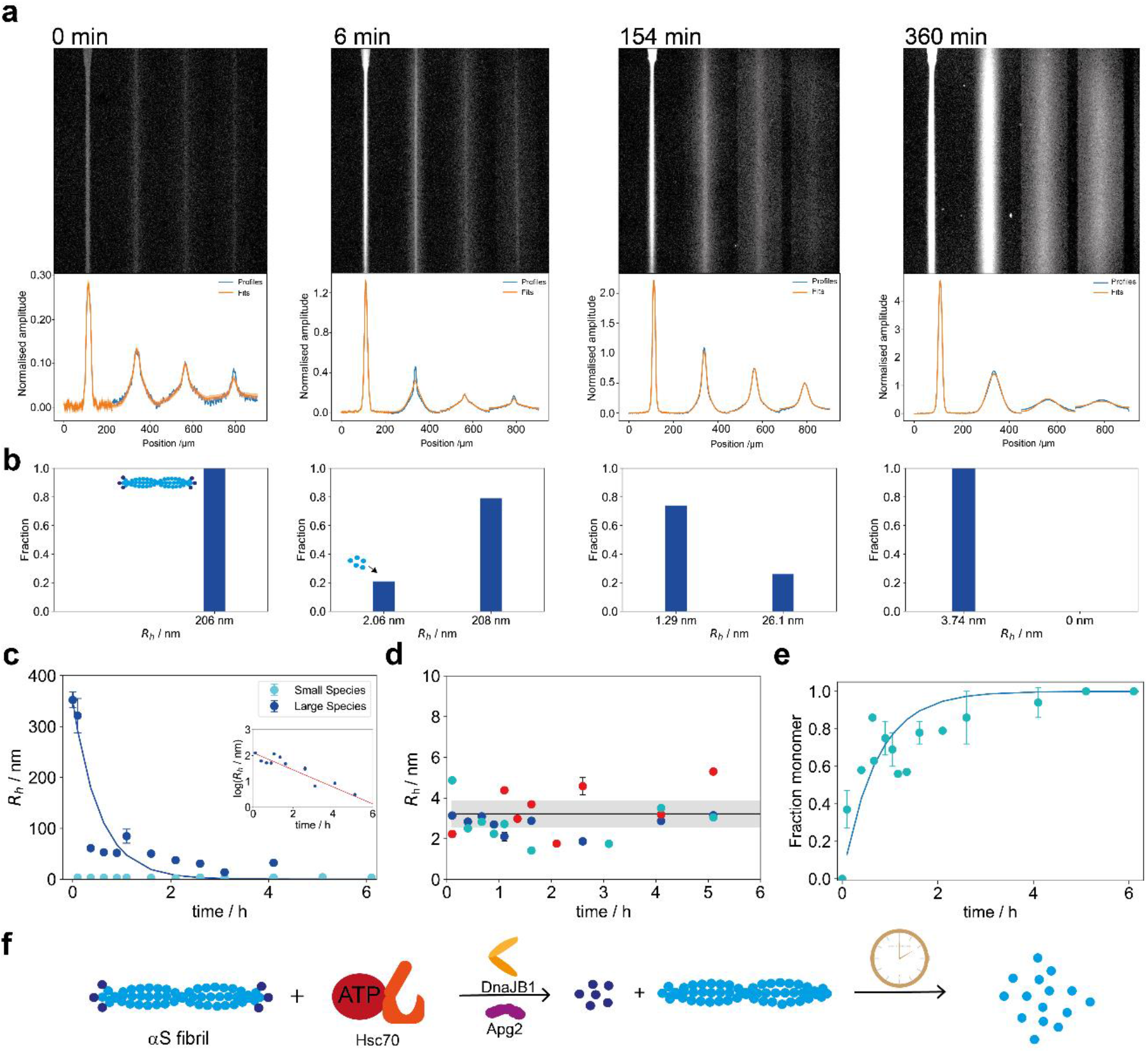
Diffusional sizing and kinetic analysis αS fibril disaggregation by the Hsc70 chaperone machinery. **(a)** Representative images of microfluidic diffusional sizing experiments at different time points and diffusion profiles obtained from image analysis during the disaggregation time course. Diffusion profiles broaden over time, indicating that a smaller species is created during the disaggregation reaction. At 4 min and 154 min, a heterogeneous population of a small, broadly diffused-out species and a larger, little diffusing species was observed, while the population was homogeneous at 0 min and 304 min. Extracted *R*_h_ values for the small and large species from diffusion profile fitting at each timepoint are reported in panel b. The experientially obtained diffusional profiles are shown in blue and the obtained fits in orange; the fit errors are reported as green bands. **(b)** Histograms showing the size distribution of the two species from image analysis. At 0 min and 304 min, *R*_h_ values of single-component fits are reported. (**c**) Evolution of *R*_h_ for small (cyan) and large species (blue) over time. The size of the larger species decayed monotonically, consistent with a single exponential fit (red line). The size of the smaller species remained constant over time. Inset: logarithmic representation of *R*_h,large_ over time. From these fits, a rate constant of *k* = 1.8 ± 0.3 · 10^–4^ s^–1^ was determined. **(d)** *R*_h_ of the small species for three independent disaggregation reactions over time (red, blue, cyan) compared to *R*_h_ of pure monomer (black line and grey region indicate the expected size and size range). This analysis shows that monomer is produced throughout the reaction and no other intermediate is generated. **(e)** Fraction of monomer over time (black), showing that the fraction of monomer increases over time, whereby less monomer is generated at later time points, consistent with Fig. 2d. Error bars in d, e and f represent the standard deviation of *n* = 3 independent experiments. **(f)** Schematic of the disaggregation reaction and the experimental assay.

In a first series of experiments, we monitored the kinetics of αS fibril disaggregation by the Hsc70 disaggregation machinery. To this end, we incubated the chaperone system consisting of Hsc70, DnaJB1, and Apg2 in presence of an ATP-regenerating system (see Methods) with preformed αS fibrils labelled with AlexaFluor488 (N-terminal amine labelling, 10% labelled monomer). Such labelled fibrils were structurally comparable to unlabelled fibrils, as shown by atomic force microscopy (Fig. S2). The sample was introduced at the centre of the microfluidic channel and the extent of diffusion towards the edges of the channel monitored as a function of channel position. Pure fibrils in the absence of the disaggregation machinery showed a monodisperse distribution and only little broadening along the microfluidic channel (Fig. 2a, 0 min), in agreement with the expected large size of αS fibrils (approx. 200-300 nm). By contrast, a clear signature of fibril disintegration became apparent in the presence of the disaggregase system as diffusion profiles broadened over the time course of the reaction.

At early time points (Fig. 2a, 4 min), fibrils were still predominant; however, a second protein species with a high degree of diffusive broadening became apparent, showing that the initiation of the disaggregation reaction is fast (i.e., within a few minutes) and that a smaller molecular weight species is being produced. At intermediate time points (Fig. 2a, 154 min), the fluorescence intensity arising from the smaller diffused out species increased relative to the larger species, while fibrils were still present. This is consistent with a scenario in which monomer units are taken off directly from fibril ends without any intermediate fragmentation steps (see also below). At late time points (Fig. 2a, 304 min), the diffusion profiles broadened significantly and yielded a monodisperse population of the smaller molecular weight species. Full disaggregation to the smaller molecular weight species was observed after ~240 min. Crucially, disaggregation required the presence of all three chaperone components and ATP. Specifically, Hsc70 alone with ATP was unable to mediate fibril disaggregation (Fig. S3).

Quantitative analysis of the diffusion profiles as a function of time revealed that the profiles are indeed best described by two diffusing species (Fig. 2c). The fraction of the larger species with an initial *R*_h_ of ~350 nm, corresponding to the size of fibrils (Fig. 2c), and decayed monotonically over time, while simultaneously the fraction of a smaller, diffused out species with a *R*_h_ of ~ 3 nm gradually increased (Fig. 2e). The smaller species, that built up in time, corresponds in size to pure monomer (Fig. 2d), demonstrating that the reconstituted Hsc70 machinery disaggregates αS fibrils into monomer units. The observation that monomer was already abundant at very early time points not only highlights the fact that the fibril-to-monomer conversion reaction is very fast (i.e., on the minutes timescale), but also strongly indicates that monomer units are removed from the fibril ends directly, as opposed to scenarios that involve intermediate fragmentation steps, in which case other species than fibrils and monomer would have been observed. No monomer is observed in the fibril-only sample (Fig. 2a, 0 min).

Further quantification revealed that the size decay of the large fibrillar species was best described with a single exponential kinetic model, yielding a rate constant of *k* = 1.8 ± 0.3 · 10^−4^ s^−1^. Crucially, the apparent size of the species converged to that of the smaller species (i.e., monomer) after ~3 hours, indicating that the disaggregation reaction has gone to completion. The single exponential behaviour was observed independent of whether experiments with full-length fibrils (Fig. 2c) or sonicated fibrils (Fig. S4 a–b) were performed. However, the reaction was slightly faster for the sonicated fibrils, indicating that the abundance of more fibril ends accelerates the reaction when fibril concentration is rate-limiting, as the Hsc70 is in excess under the disaggregation reaction conditions, with a significant portion of Hsc70 remaining unbound during the reaction cycle (Fig. S5). A single exponential behaviour is consistent with a pseudo-first order kinetic model (Fig. 2c) and thus supportive of a one-step disaggregation reaction mechanism, showing that the monomer is taken off the fibrils directly, likely from fibrils ends, without previous fragmentation of the fibril. This is also consistent with the gradual increase of monomer concentration over time and the observation that monomer is generated almost immediately after addition of the chaperones (Fig. 2a).

### Single Round Disaggregation Experiments

To further substantiate the above findings, the outcome of a single round of disaggregation was measured and the products characterised. For this purpose, the reaction mixture was incubated with DnaJB1, Hsc70, αS fibrils and the ATP regeneration system for 5 min to ensure binding of the Hsc70 to the fibrils. Subsequently, a 200-fold molar excess of the Hsc70 binding peptide GSGNRLLLTG^28–30^ was added and substrate release triggered with Apg2 (Fig. 3a). Due to the presence of Hsc70 binding peptide, the substrate was no longer able to rebind to Hsc70. Addition of the Hsc70 binding peptide to an ongoing disaggregation stopped the disaggregation reaction immediately (Fig. S6a), and no disaggregation was observed upon addition of the peptide before adding Hsc70 (Fig. S6). This assay thus allowed monitoring the outcome of only a single round of disaggregation. As shown in Fig. 3a, two species were observed: a smaller species with a radius of *R*_h_ = 2.71 ± 0.08 nm, corresponding to monomer, and a larger species with radius of *R*_h_ = 233 ± 22 nm, corresponding to the fibril size (c.f. *R*_h_ = 235 ± 28 nm) prior to disaggregation. Due to the conserved fibril length, our data further supports the notion that monomer is taken off the fibril ends rather than from the fibril core, given that no other species of intermediate size between fibril and monomer is observed. In case of a fragmentation mechanism, in contrast, fibrillar species of smaller length would be observed. The same behaviour was found when the reaction was carried out with equimolar concentrations of ATP and an excess of the slowly hydrolysable ATP analogue ATP-γ-S (Fig. S6c).

**Figure 3:**
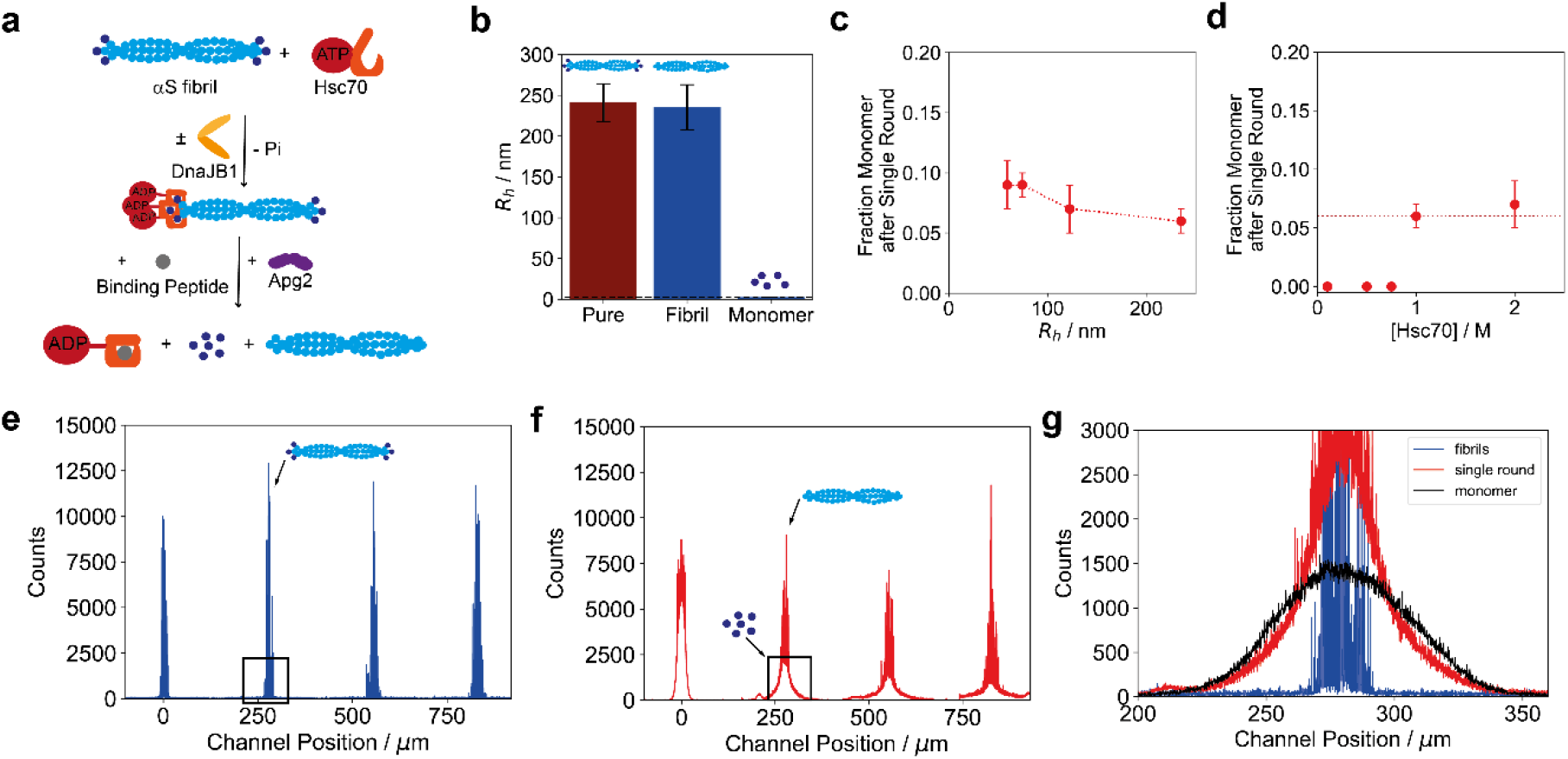
Single round disaggregation experiments. **(a)** To measure the outcome of a single disaggregation round, Hsc70, DnaJB1 and the αS fibrils (2 μM monomer equivalent) were incubated. Subsequently, a 200-fold molar excess of Hsc70 binding peptide GSGNRLLLTG was added, leading to the blockage of the Hsc70 binding site and preventing rebinding, thereby allowing us to measure the outcome of a single disaggregation round on αS fibrils. **(b)** Outcome of a single disaggregation round. Two species were detectable, the larger of which had a similar size to the initial fibril, the second of which had the size of monomer. This suggests that monomer is taken off the fibril ends directly, as, otherwise, the larger species would be significantly shorter than the initial fibril length. **(c)** Fraction of monomer generated as a function of the fibril length. Shorter fibrils lead to more monomer release than longer fibrils, whereby this effect plateaus for fibrils longer than 100 nm. **(d)** Fraction of monomer measured at different Hsc70 concentrations. No disaggregation took place for Hsc70 concentrations below 1 μM, whereas there was no significant difference in the monomer fraction released from fibrils at higher Hsc70 concentrations. This suggests that there is a critical Hsc70 concentration required for efficient disaggregation. Error bars in b, c and d represent the standard deviation of *n* = 3 independent experiments.

Interestingly, the fraction of monomer generated during the disaggregation reaction increased with shorter fibril lengths under constant Hsc70 concentrations (Fig. 3c). Shorter fibrils are consistent with an increased concentration of fibril ends; therefore, if monomer units are removed from the fibril ends, this behaviour is expected, while no effect of a decreasing fibril length distribution would be expected in case of a fragmentation mechanism. Furthermore, this effect levels off for fibril lengths above 100 nm, indicative of a critical ratio between Hsc70 concentration and the number of fibril ends for effective disaggregation. To validate this hypothesis, single round experiments were performed with fibrils of identical length distributions but with varying concentration of the Hsc70 chaperone (Fig. 3d). This experiment revealed that, at an Hsc70 concentration below 1 μM, no disaggregation is observable. This result is interesting, as it suggests a cooperative action or clustering of multiple Hsc70 molecules at individual fibril ends.

To further explore the molecular details of the dissociation mechanism, we monitored the outcome of a single round of disaggregation using confocal microscopy. Through integration of single-molecule scanning optics with microchip diffusional sizing, individual fibrils can be detected directly while the presence of monomeric αS generated during single disaggregation reactions can be probed and quantified simultaneously as well. We first performed experiments on pure fibrils (i.e., in the absence of the disaggregation machinery) (Fig. 3e). Diffusion profiles showed large bursts located at the centre of the channel and profile shapes that do hardly broaden along the channel, as expected for large amyloid fibrils containing of a large number of labelled monomer units. Fitting of the profile yielded a hydrodynamic radius of *R*_h_ = 219.9 ± 11.2 nm consistent with the diffusional sizing experiments in epifluorescence mode and the expected size of fibrils. Strikingly, after a single round of disaggregation, a diffused out monomeric population with a hydrodynamic radius *R*_h_ = 2.46 ± 0.01 nm appeared in addition to the fibril bursts located at the centre of the channel; their hydrodynamic radius of *R*_h_ = 204.3 ± 3.1 nm was conserved compared to the fibril-only experiment. Comparing the diffusion profile of the monomer population obtained in the single round disaggregation experiment with a concentration series performed on labelled αS monomer (Fig. S7) revealed that approximately 100 nM of monomeric protein is present after the single disaggregation round. This suggests that ~5 % of the αS population is monomeric after a single round of disaggregation. Importantly, no intermediate species between monomer and fibrils were observed. Together, these findings support the notion that monomer is taken directly off the fibril ends.

### Interactions between Chaperones their Binding to Fibrils

Having established that all components of the chaperone system are necessary to successfully disaggregate αS fibrils (Fig. S5), we next focused on the interactions between these chaperones as well as between the chaperones and the fibrils in order to gain further mechanistic insights. Upon binding of a labelled chaperone to an unlabelled chaperone or fibril, a decrease in its molecular diffusivity concomitant with an increase in its effective molecular weight and size upon binding is expected, thereby allowing us to determine the binding affinity of the respective interactions^16^.

First, we performed experiments involving Hsc70 in the presence of ATP, ADP, and its non-hydrolysable analogue ATP-γ-S. As shown in Fig. 4a, Hsc70 did undergo a conformational change upon binding of ATP or ATP-γ-S, as reflected in a decrease in its hydrodynamic radius. Conversely, Hsc70 has a similar conformation in the ADP bound state relative to apo-Hsc70. These observations are consistent with structural analyses of the bacterial Hsp70 homolog showing that in the ATP state the hydrophobic interdomain linker and the α-helical lid of the SBD are associated with the NBD, and the SBD is in an open conformation, whereas the SBD and NBD are loosely associated in the ADP state^43–47^.

**Figure 4:**
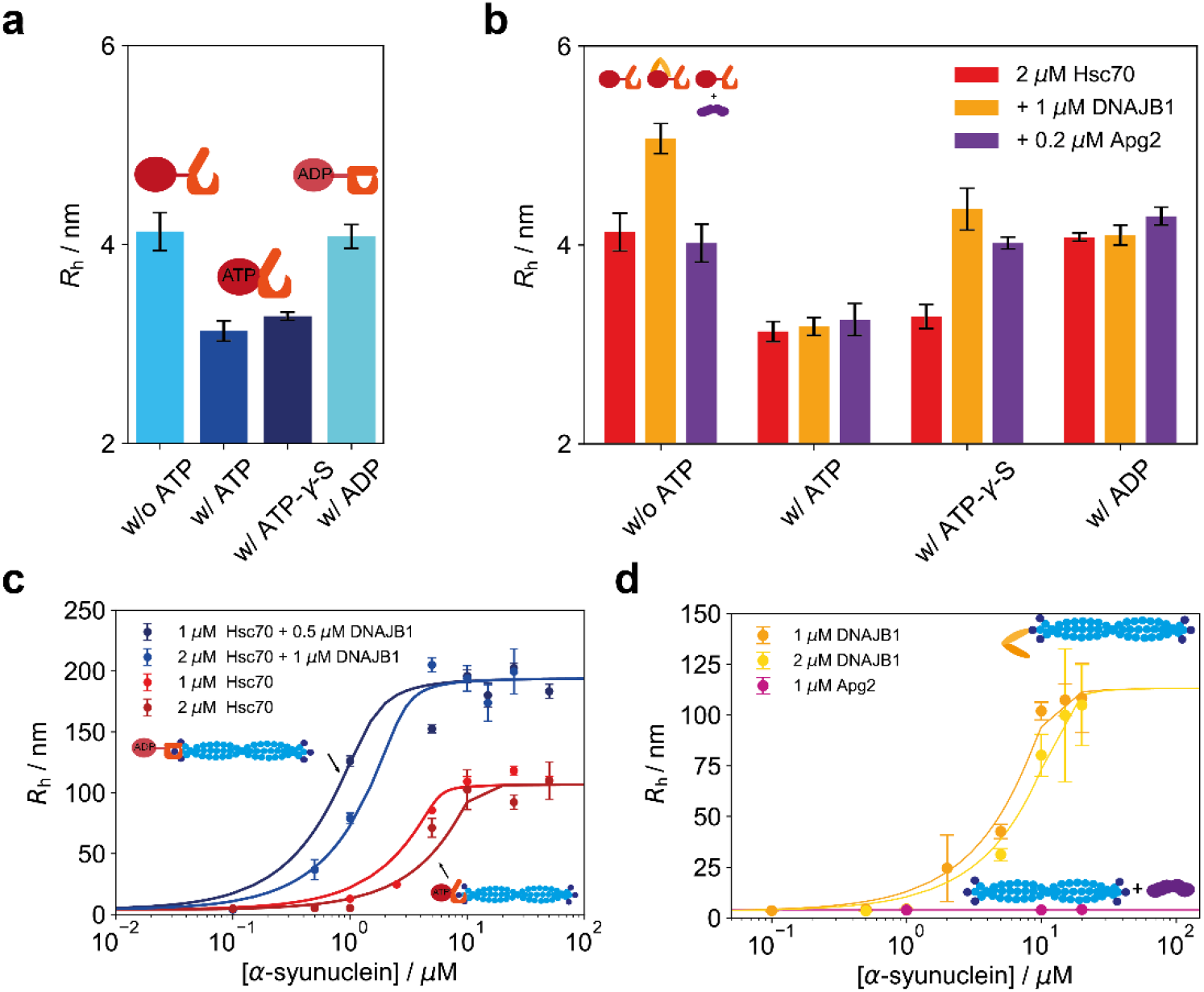
Characterisation of the binding interaction between chaperones and αS fibrils. **(a)** Conformational change of fluorescently labelled Hsc70 upon ATP binding, as envisioned by the change in hydrodynamic radius. When binding to ADP, the effect is reverted again. **(b)** Binding interaction between fluorescently labelled Hsc70 and its co-chaperones. Binding was observed between Hsc70 and DnaJB1, leading to an increase in hydrodynamic radius. DNAJB1 can bind in absence of ATP and in presence of ATP-γ-S, whilst it dissociates from Hsc70 in the ATP bound state due to fast hydrolysis of ATP. Error bars are standard deviations from triplicate measurements. **(c)** Binding of Hsc70 to αS fibrils. Hsc70 shows a binding affinity of *K*_d_ = 138.07 ± 22.89 nM in the presence of ATP and a tighter affinity *K*_d_ = 83.95 ± 8.1 nM in the presence of DnaJB1, which triggers hydrolysis of ATP to ADP, thereby leading to lid closure and stronger binding. Error bars are standard deviations from triplicate measurements. **(d)** Binding of DnaJB1 and Apg2 to αS fibrils. Apg2 did not bind to αS fibrils. DnaJB1 bound to αS fibrils with an affinity of *K*_d_ = 246.1 ± 28.1 nM. Error bars are standard deviations from triplicate measurements. The different plateau levels in b and c are caused by different batches of αS fibrils; the concentrations of αS are given with respect to monomer equivalents.

Next, we performed sizing experiments of Hsc70 in the presence of DnaJB1 or Apg2 (Fig. 4b). Hsc70 bound to DnaJB1 in the absence of ATP with an affinity of *K*_d_ = 46.0 ± 13.5 nM (Fig. S8c), but Hsc70 did not bind to DnaJB1 in the presence of ATP or ADP; however, binding was detectable in presence of ATP-γ-S. DnaJB1 is known to accelerate the hydrolysis of ATP by Hsc70, and is likely to dissociate from ADP-bound Hsc70^12^. This is consistent with the results obtained here. Evidently, the ATP-state of Hsc70 is short-lived in the presence of DnaJB1, precluding detection of binding in the fluidics system. However, a significant size increase of Hsc70 due to DnaJB1 binding was observed in the presence of the non-hydrolysable ATP analogue ATP-γ-S, which prolongs the ATP-state of Hsc70. Similarly, Hsc70 showed only weak binding to Apg2, consistent with a transient interaction during nucleotide exchange. The size Apg2 was large (*R*_h_ = 4.46 ± 0.20 nm) (Fig. S8b), which may contribute to its effectiveness in the disaggregation reaction in an entropic pulling mechanism^36–38^.

Subsequently, we investigated the binding between Hsc70 and fibrils. As shown in Fig. 4b in presence of hydrolysable ATP, a binding affinity of *K*_d_ = 138.07 ± 22.89 nM is observed, which is increased to *K*_d_ = 83.95 ± 8.1 nM in the presence of DnaJB1, consistent with DnaJB1 accelerating the hydrolysis of ATP to ADP. Furthermore, stoichiometry analysis of Hsc70 binding to fibrils yielded one Hsc70 chaperone molecule per 5.3 ± 0.6 monomeric units of αS within the fibril. Notably, this stoichiometric ratio represents an average value that is consistent either with a scenario of one Hsc70 binding site every 5–6 αS monomer units within the fibril, or a scenario where clustering of Hsc70 occurs at fewer binding sites on the fibrils. Crucially, our kinetic analysis above reveals that monomer is taken from the fibril ends. Hence, given that we observed no disaggregation at decreased Hsc70 concentrations (Fig. 3c), our results strongly support the scenario that clustering of Hsc70 at fibril ends is likely a crucial feature of the disaggregation mechanism.

Lastly, we investigated the binding between αS fibrils and the co-chaperones. While no binding of Apg2 alone to the fibrils was detectable, DnaJB1 bound the fibrils with an affinity of *K*_d_ = 246.1 ± 28.1 nM (Fig. 5c). This supports that DnaJB1 recruits Hsc70 to the fibrils, as the affinity of DnaJB1 to the fibrils is significantly lower than that of Hsc70.

**Figure 5:**
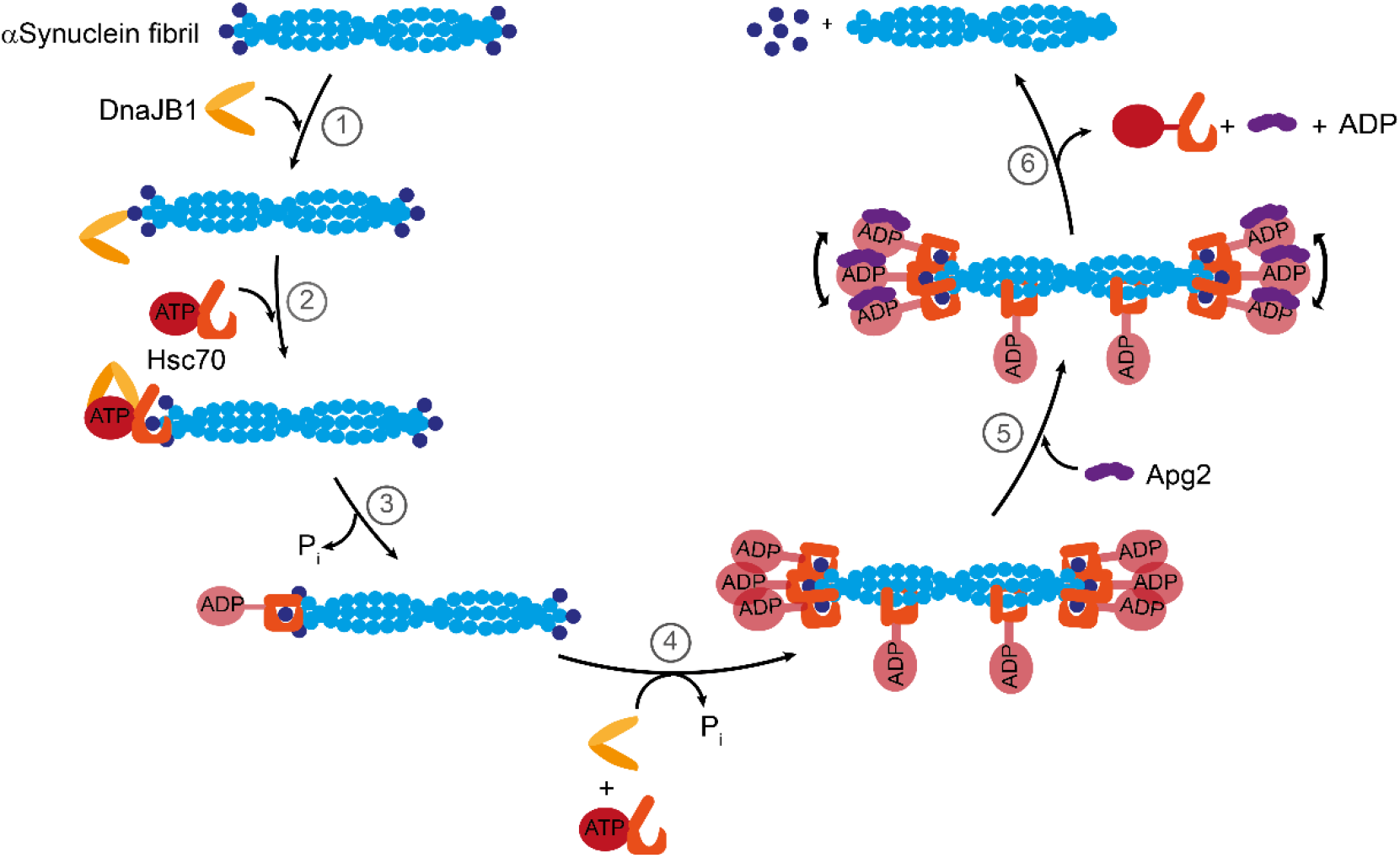
Emerging principle of Hsc70 disaggregase activity. Model for Hsc70-mediated protein disaggregation of αS fibrils. This disaggregation mechanism comprises six steps: step 1, DnaJB1 binding to αS fibrils; step 2, Hsc70 loading; step 3, hydrolysis of ATP to ADP and dissociation of DNAJB1; step 4: further Hsc70 loading resulting in clustering of Hsc70 at fibril ends; step 5, Apg2 binding to Hsc70; step 6, dissociation of αS monomer.

## Discussion

Hsc70 together with its co-chaperones DnaJB1 and Apg2 (Hsp110) have been shown to disaggregate αS fibrils^18^. By bringing together microfluidic measurements with chemical kinetics and thermodynamic analysis, we have investigated this process here in a quantitative manner and uncovered the mechanistic determinants of this process. We have found that, during the disaggregation reaction, only two species are significantly populated, namely larger, fibrillar species and αS monomer units released from the fibrils. The time dependent size decrease of the larger species is consistent with pseudo-first order kinetics. This observation is further supported by the fact that there is an excess of unbound Hsc70 in the disaggregation reaction mixture (i.e., the concentration of fibrils is limiting). Thus, αS monomers are taken off the fibrils directly. Indeed, pure monomer is abundant almost immediately after the addition of the chaperones, and the monomer fraction increases linearly. An advantage of microfluidic diffusional sizing is that the end product of the disaggregation reaction, monomeric αS, can be detected, which is not ThT active and thus not observable in traditional disaggregation assays. Interestingly, our results indicate that the disaggregation reaction is independent of the initial fibril lengths (i.e., whether full-lengths fibrils or sonicated fibrils were used), while the rate is slightly accelerated for shorter fibrils. As a single round of chaperone action produces only monomer and fibrils, this further suggests that monomer is taken off the fibril ends. Dissociation of monomer from within the fibrils would have resulted in a substantial decrease in fibril length due to induced fragmentation. Future directions will include investigating the effect of different fibril polymorphs and αS variants on the disaggregation kinetics.

Our analysis of the disaggregation kinetics and the interactions between the chaperone components and their interactions with αS fibrils, together with previous reports^7^, leads to a picture of chaperone function as shown in Figure 5 and suggests that the disaggregation reaction can be divided into the following six steps: DnaJB1 binds first to the fibrils (step 1) and recruits Hsc70 in the ATP state. ATP hydrolysis on Hsc70, accelerated by DnaJB1, results in tight binding of Hsc70 to fibrils (step 2), subsequent hydrolysis of ATP to ADP (step 3), and clustering of Hsc70 on the fibril ends (step 4, see below). Disaggregation resulting in αS monomer production then results upon addition of Apg2 (step 5/6). As Apg2 alone does not detectably interact with the fibrils, it may act solely as a NEF, although an interaction with the fibril substrate in the presence of Hsc70 cannot be ruled out. The large size of Apg2 may facilitate disaggregation according to the model of entropic pulling^36–38^. Further studies should be directed to understand the role of the NEFs, especially of the relevance of their size in the interaction. Our data suggest further that disaggregation requires clustering of Hsc70 molecules, with repulsive forces between αS-bound Hsc70 molecules inducing monomer dissociation from fibril ends (step 5). This could be mechanistically similar to the observed conformational expansion of non-native proteins by the binding of multiple Hsp70 to sites within the same polypeptide chain^30,48^. Alternatively, since αS monomer dissociation from the fibril critically depends on Apg2, we speculate that binding of Apg2 to clustered Hsc70 molecules induces the steric repulsion that drives disaggregation (Fig. 5). Further directions to extend on our studies should include investigation of these phenomena with various other substrates, including the amyloid-β peptide or Tau protein, key players in the onset and progression of Alzheimer’s disease.

In conclusion, we present here a comprehensive kinetic and thermodynamic description of a complex protein quality control mechanism which is likely to be crucial in the clearance of amyloidogenic deposits of the protein αS. We found that, in the disaggregation reaction of αS fibrils, monomer is taken directly from the fibril ends and that the Hsc70 chaperone system assembles in clusters at the fibril ends. This process follows single exponential kinetics and is highly effective, such that it leads to complete disaggregation of amyloid fibrils, thereby likely contributing to prevention of Parkinson’s diseases.

## Methods

### Materials

All chemicals were purchased in the highest purity available. Hydroxyethyl piperazineethanesulfonic acid (HEPES), potassium hydroxide (KOH), potassium chloride (KCl), dithiothreitol (DTT), and Tween20 were purchased from Sigma Aldrich (St Louis, MO, USA) and were of analytical grade. Pyruvate kinase (10109045001) from rabbit muscle was purchased from Roche (Basel, Switzerland). Poly-(dimethylsiloxane) (PDMS) and curing agent were purchased from Momentive (Techsil, Bidford-on-Avon, UK). For all microfluidic experiments, the buffer used contained 50 mM HEPES-KOH (pH 7.5), 50 mM KCl, 5 mM MgCl_2_, 2 mM DTT; co-flow buffer was additionally supplied with 0.01% Tween20.

### Expression and purification of molecular chaperones and αS monomer

Recombinant human wildtype αS was purified similarly as described previsouly^49^. In brief, *Escherichia coli* BL21(DE3) cells were transformed with pT7-7 αS and cultured in lysogenic broth (LB) medium. Protein expression was induced by 1 mM isopropyl β-D-1-thiogalactopyranoside (IPTG) for 4 h at 37°C. Bacteria were harvested and pellets were lysed in 10 mM Tris–HCl (pH 8.0), 1 mM EDTA, 1 mM phenylmethylsulfonyl fluoride (PMSF). The lysate was sonicated for 5 min and boiled subsequently for 15 min, followed by centrifugation. The supernatant was subjected to streptomycin sulphate and ammonium sulphate precipitation steps as described. The ammonium sulphate pellet formed after centrifugation at 5’200 g for 30 min was dissolved in 50 mM Tris–HCl (pH 7.5), 150 mM KCl and subjected to size exclusion chromatography (SEC) on a Superdex 200 column (GE Healthcare, Chalfont St Giles, UK). αS fibrils were generated by shaking a mixture containing 10% labelled and 90% unlabelled monomer at 37°C and 200 rpm for 4 days. Fibrils were sonicated (cycle 0.3, power 10%, 90 seconds) using a Sonopuls ultrasonic homogenizer (Bandelin, Nänikon, Switzerland).

Human Hsc70, DnaJB1 and Apg2 were expressed in *E. coli* BL21(DE3) as fusion proteins with protease-cleavable His_6-_ or His_6-_Smt3 tags and purified by tandem Ni-affinity chromatography with intermittent protease cleavage similar as described previously^50^.

Hsc70 was expressed from the plasmid pProEx-HtA Hsc70. Cells were grown in LB medium at 37°C to an OD_600_ ~0.5 and induced with 0.5 mM IPTG for 18 h at 21°C. Cells were lysed by ultrasonication in 50 mM HEPES–KOH (pH 8.0), 10 mM KCl, 5 mM MgCl_2_ (buffer A) containing 0.8 mg mL^−1^ lysozyme at 4 °C. The supernatant after centrifugation at 125,000 g for 45 min was applied to a Ni-NTA column (GE Healthcare, Chalfont St Giles, UK) equilibrated in buffer A. The column was washed with a step gradient of buffer A containing increasing amounts of imidazole (20/250/1000 mM). The bound protein was eluted with buffer A containing 250 mM imidazole. This was followed by cleavage of the His_6_-moiety at 4°C with His_6_-tobacco etch virus (TEV) protease during 45 h. After transfer into buffer A using a desalting column, the material was passed over the Ni-NTA column and the flow-through collected. Next, Hsc70 was purified by anion exchange chromatography on MonoQ (GE Healthcare, UK) in the same buffer system using a linear salt gradient (0–700 mM KCl) in 50 mM HEPES–KOH (pH 8.0), 5 mM MgCl_2_. Finally, the Hsc70-containing fractions were subjected to SEC on Superdex 200 column (GE Healthcare, Chalfont St Giles, UK) in 50 mM HEPES–KOH (pH 8.0), 150 mM KCl, 5 mM MgCl_2_ and 5% glycerol.

DnaJB1 and Apg2 were expressed from the plasmids pCA528-DnaJB1 and pCA528-HspA4, respectively^17^. Cells were grown in LB medium at 37°C to an OD_600_ ~0.5 and induced with 0.5 mM IPTG for 5.5 h at 30°C. Cells were lysed with an Emulsiflex (Avestin, Ottawa, Canada) cell disruptor in 50 mM HEPES–KOH (pH 7.4), 10 mM KCl, 5 mM MgCl_2_ (buffer B) containing 2 mM PMSF and Complete protease inhibitor cocktail (Roche). After centrifugation, the supernatant was applied to a Ni-NTA column equilibrated in buffer B. After washing with buffer B, the bound protein was eluted with buffer B containing 250 mM imidazole. The His_6_-Smt3 moiety was cleaved with PEN2 protease (MPIB Core facility) at 4°C in the presence of 1 mM dithiothreitol (DTT). After buffer exchange, the mixture was passed over the Ni-NTA column and the flow-through collected. SEC on Sephacryl S-200 (GE Healthcare, UK) in 50 mM Tris–HCl (pH 8.0), 5 mM MgCl_2_ and 150 mM KCl (buffer C) was the final purification step for Apg2. DnaJB1 was further purified by SEC on Sephacryl S-100 in buffer C containing 5% glycerol, and by cation exchange chromatography on Source 30S (GE Healthcare), wherein the elution was carried out with a linear salt gradient (0–400 mM NaCl) in 50 mM Tris–HCl (pH 7.5).

### Microfluidic diffusional sizing

A scheme of the chip design is shown in Fig. 1a. Fabrication and operation of the microfluidic devices for microfluidic diffusional sizing have been described previously^41,51,52^. Briefly, the microfluidic devices were fabricated in PDMS by standard soft-lithography techniques and bonded onto a glass coverslip after activation with oxygen plasma. Sample loading from reservoirs connected to the respective inlets and control of flow rate was achieved by applying a negative pressure at the outlet using a glass syringe (Hamilton, Bonaduz, Switzerland) and a syringe pump (neMESYS, Cetoni GmbH, Korbussen, Germany). A custom-built inverted epifluorescence microscope equipped with a charge-coupled-device camera (Prime 95B, Photometrics, Tucson, AZ, USA) and brightfield LED light sources (Thorlabs, Newton, NJ, USA) was used to record the images. Images were typically taken at flow rates 20 μL/h, 60 μL/h, and 100 μL/h, and lateral diffusion profiles were recorded at 4 different positions along the microfluidic channels.

Diffusion profiles extracted from fluorescence images using a custom-written analysis software were fitted by numerical model simulations by solving the diffusion–advection equations for mass transport under flow^41^. For evaluation of the disaggregation time courses, we assumed two species representing the fibril and the dissociated monomers. For the thermodynamic evaluation and sizing of pure species, we fitted the diffusion profiles with one species only to determine the average size of the bound and unbound chaperone.

### Kinetic measurements

The reaction mixture for disaggregation measurements contained 2 μM αS fibrils, 2 μM Hsc70, 1 μM DnaJB1, 0.2 μM Apg2, 5 mM 2-phosphoenolpyruvate, 0.05 mg/mL pyruvate kinase, and 5 mM ATP in 50 mM HEPES-KOH (pH 7.5), 50 mM KCl, 5 mM MgCl_2_, 2 mM DTT. This mixture was incubated at 300 rpm and 30°C. At different time points, 8 μL aliquots were taken out, injected into the microfluidic chip and images were acquired as described above. The kinetics were fitted with a pseudo-first order kinetic model, with the following rate law, as the concentration of fibrils, m, evolves with time as follows:

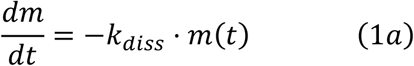

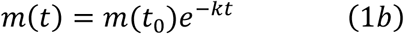

Therefore, the hydrodynamic radius at time t is described with

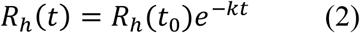

Likewise, the rate of the increase concentration of monomer, M, increases as

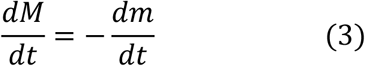

or, in its integrated and normalised form, the fraction of monomer, *F*_M_, becomes

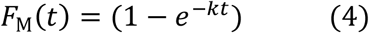

### Single round disaggregation experiments

The reaction mixture for disaggregation measurements contained 2 μM αS fibrils, 2 μM Hsc70, 1 μM DnaJB1, 5 mM 2-phosphoenolpyruvate, 0.05 mg/mL pyruvate kinase, and 5 mM ATP in 50 mM HEPES-KOH (pH 7.5), 50 mM KCl, 5 mM MgCl_2_, 2 mM DTT. This mixture was incubated at 300 rpm and 30°C for 5 min. Subsequently, 0.2 μM Apg2 and 400 μM Hsc70 binding peptide GSGNRLLLTG^28–30^ were added. This peptide can bind to Hsc70, thereby preventing substrate rebinding and, thus, terminating the disaggregation.

Diffusional sizing experiments involving confocal microscopy were done on a custom-built laser confocal microscopy setup. Briefly, the microscope is equipped with a 488-nm laser line (Cobolt 06-MLD, Hübner Photonics, Derby, UK) and a single-photon counting avalanche photo diode (SPCM-14, PerkinElmer, Seer Green, UK) for subsequent detection of emitted fluorescence photons. Further details of the optical unit have been described previously^53^. Diffusion profile recording was done by continuously moving the confocal observation volume through the centre four channels of the microfluidic device.

### Thermodynamic characterisation

For binding experiments, samples were prepared in typically 30 μL total volume, using the same working concentrations of the interacting partners considered as in the disaggregation time course described above under the same buffer conditions. The protocol for the equilibrium binding curves was adapted from previous reports^16^. The concentration of one of the interacting molecules was varied between 0.1 μM and 10 μM accordingly, while the labelled component was held at a constant concentration equal to the working concentration discussed previously. The samples were typically incubated for 30 min and then measured in triplicates in three independent channels at three flow rates.

## Acknowledgements

The research presented in this manuscript has received funding from the European Research Council (ERC) under the European Union’s Seventh Framework Programme (FP7/2007-2013) through the ERC grant PhysProt (agreement no. 337969) and under European Union’s Horizon 2020 research and innovation programme (ETN grant 674979-NANOTRANS) (MMS, TWH, EA, QAEP, TPJK), the Marie Skłodowska-Curie grant (agreement no. 749370) (SG) and Marie Skłodowska-Curie grant MicroSPARK (agreement no. 841466) (GK). This research has been supported by the Centre for Misfolding Diseases (CMD, TPJK), the Frances and Augustus Newman Foundation (TWH, TPJK), the Welcome Trust (CMD, TPJK), the Engineering and Physical Sciences Research Council (EPSRC) (TPJK), St John’s College Cambridge, UK (MMS, CMD, TPJK), the Oppenheimer Early Career Research Fellowship (TWH), the Herchel Smith Funds of the University of Cambridge (GK), and the Wolfson College Junior Research Fellowship (GK). We are grateful to Romy Lange and Nadine Wischnewski for their assistance with protein expression. Plasmids pCA528-HspA4 and pCA528-DnaJB1 were kind gifts from Bernd Bukau, University of Heidelberg.

## Author Contributions

FUH, TPJK, CMD, AB, SG, GK, TWH and MMS designed the study. MMS, TWH, SG, EA, GK, AMM and FSR performed the experiments. SG, GK, AB, CMD, FUH and TPJK provided materials and methods. MMS, TWH, EA, GK, QAEP, FUH and TPJK analysed the data. MMS, TWH, GK, AB, FUH, TPJK wrote the paper. All authors discussed the results and commented on the manuscript.

## Data availability

The raw data and analysis code underlying this study will be made available upon reasonable request.

## Supplementary Information

Supplementary Figures available online.

