## Supporting Information for "The Hsc70 Disaggregation Machinery Removes Monomer Units Directly from α-Synuclein Fibril Ends"

[†] Deceased (September 2019)

### Supplementary Information

### Supplementary Figures

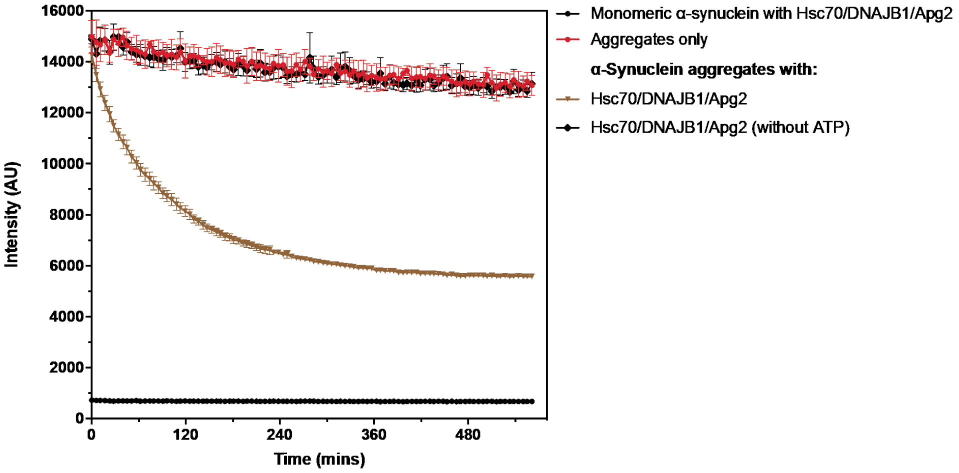


**Figure S1 :** Disaggregation Kinetics of $\alpha$-synuclein by the Hsc70-DNAJB1-Apg2 triade system, followed by ThT. As it can be see, the ThT fluorescence is decreasing over time if all chaperones are present, but it is not possible to say what the end-product of the disaggregation reaction is.

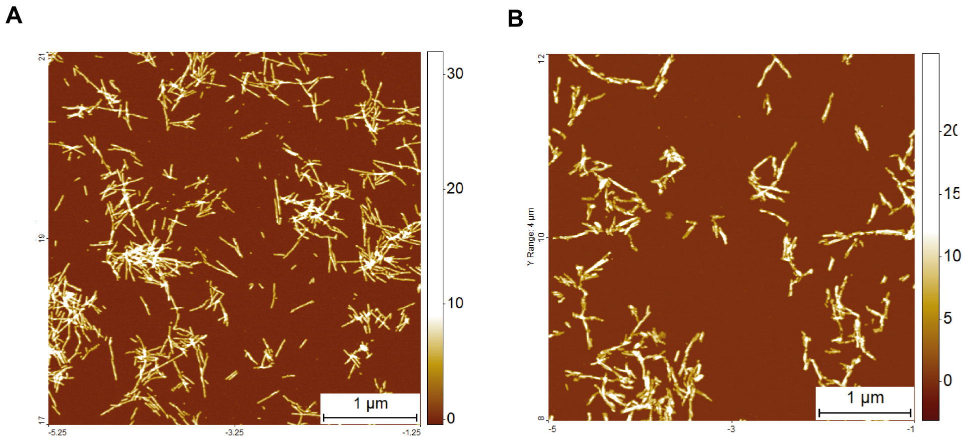


**Figure S2 :** AFM imaging to compare labelled and unlabelled fibrils. (**A**) labelled and (**B**) unlabelled fibrils were imaged by atomic force microscopy (AFM). Both labelled and unlabelled contain amyloid fibrils of similar length.

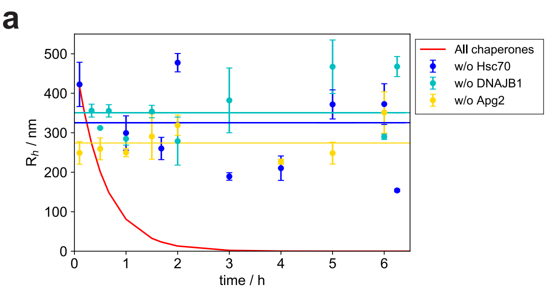


**Figure S3 :** Dependence of disaggregation on co-chaperones and ATP. (**a**) Time courses in absence of the chaperone Hsc70 (blue), as well as the co-chaperones DNAJB1 (cyan) and Apg2 (yellow). No decrease in hydrodynamic radius, $R_{\mathrm{h}}$, is observable after 6 hours, whilst a size decrease is observable in presence of all chaperones (red). This shows that the disaggregation does not occur in absence of any of these components, indicating the necessity for all of them to be present. (**b**) Time courses in absence of ATP. Also, in this case, no size decrease is observable, showing ATP dependence of the reaction.

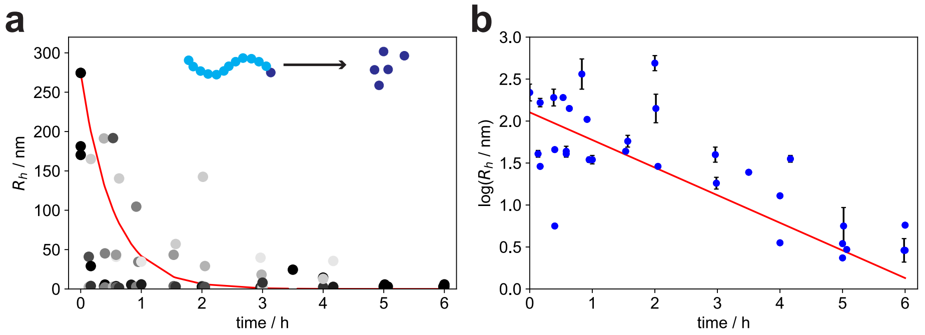


**Figure S4 :** Disaggregation of sonicated $\alpha$-syn fibrils. (**a**) Triplicate measurement of the size population of the two species over time. The size of the larger species decays, whereby the decay follows a single exponential fit (red). The size of the smaller species appearing has a conserved size (see Fig. 2d). (b) Kinetic fits of $\log(R_{\mathrm{h}})$ vs. time, according to the kinetic model described in equation 1. From these fits, a dissociation constant $k = 2.2 \cdot 10^{-4} \pm 0.2 \cdot 10^{-4}$ s$^{-1}$ was determined.

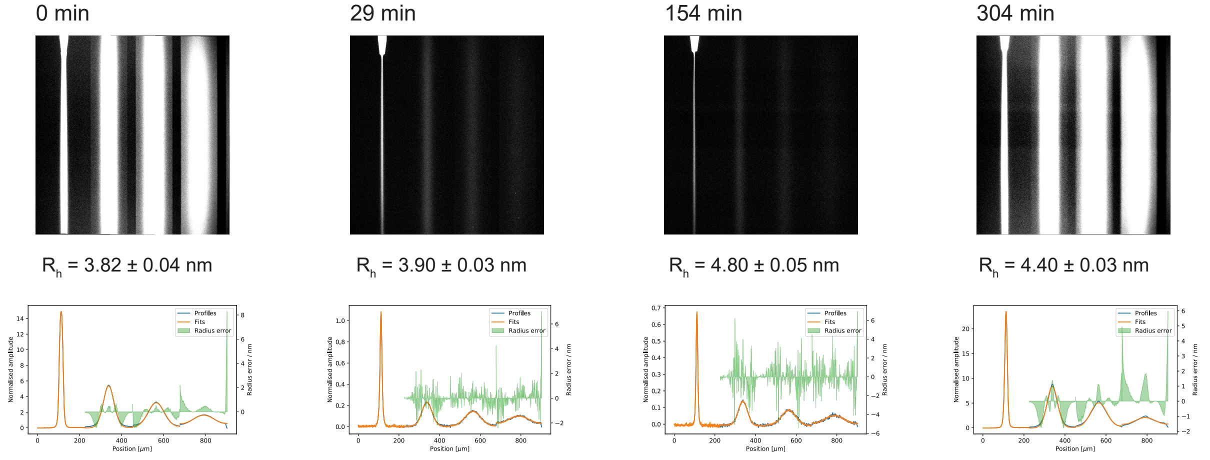


**Figure S5 :** Hydrodynamic radius of Hsc70 at different times during the disaggregation. The small size ($R_{\mathrm{h}} \approx 3.9$ nm) indicates that the majority of the Hsc70 remains in an unbound with only little Hsc70 in the bound state, indicating an excess of Hsc70 chaperone. Time point at t=0 is pure Hsc70 before mixing with $\alpha$-synuclein fibrils.

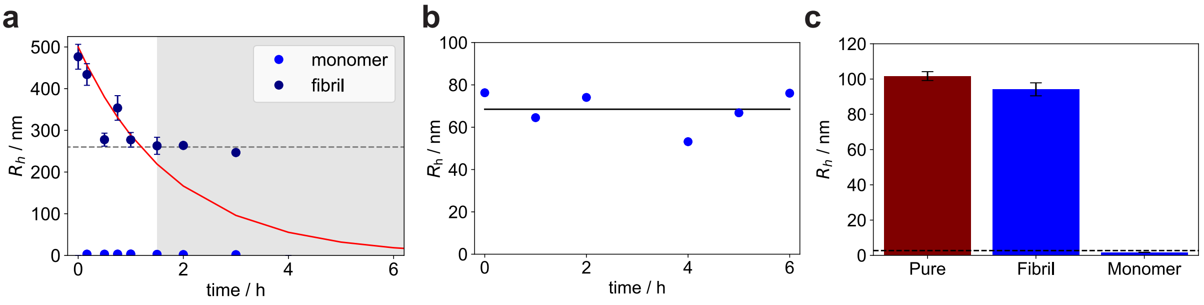


**Figure S6 :** Inhibition/Quenching of the $\alpha$S disaggregation by Hsc70 binding peptides and slowly hydrolysable ATP analogue. (**a**) Disaggregation time course was followed for 60 minutes. Then, the Hsc70 binding peptide was added, and the disaggregation time course was followed for another 120 minutes. Thereby, it can be seen that the disaggregation was fully stopped by the presence of the Hsc70 binding peptide. (**b**) Disaggregation time course with initial addition of Hsc70 binding peptide. Also in this case, the hydrodynamic radius is conserved over 6 hours, showing that the disaggregation does not proceed in presence of the Hsc70 binding peptide. (**c**) Single Round experiment with the slowly hydrolysable ATP analogue ATP-$\gamma$-S, showing the occurrence of monomer and fibril after a single round of disaggregation.

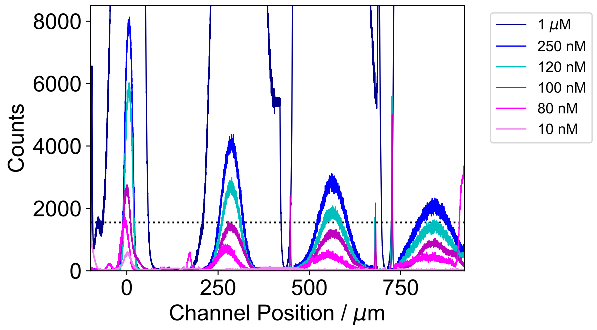


**Figure S7 :** Fluorescence Intensity of $\alpha$-synuclein monomer at different concentrations, used as calibration for the data in Fig. 3g.

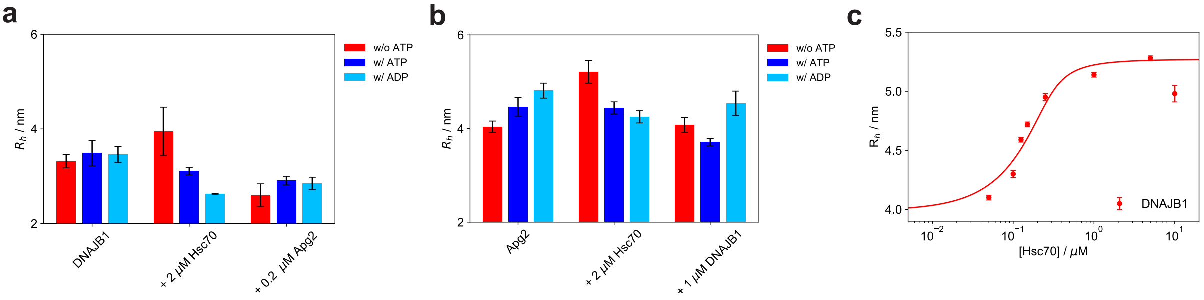


**Figure S8 :** Binding of Co-Chaperones. Binding of (**a**) labelled DNAJB1 and (**b**) labelled Apg2 to Hsc70 with different ATP/ADP conditions, co-chaperone, yielding results consistent with Fig. 4a for labelled Hsc70. (**c**) Binding curve for the interaction between Hsc70 and DNAJB1 ($K_\mathrm{d} = 46.0 \pm 13.5 nM$).
